# Minimum conductance in leaves—cuticle, leaky stomata, or water vapor saturation?

**DOI:** 10.1101/2020.05.28.120634

**Authors:** Jun Tominaga, Joseph R. Stinziano, David T. Hanson

**Affiliations:** Department of Biology, The University of New Mexico, Albuquerque, NM 87104, USA; Graduate School of Integrated Sciences for Life, Hiroshima University, Hiroshima 739-8526, Japan

**Keywords:** Cuticle conductance, Drought, Gas exchange, Model, Nighttime, Photosynthesis, Stomatal leakiness, Unsaturation

## Abstract

- Minimum conductance (*g*_*w,min*_) in leaves is important for water relations in land plants. Yet, its regulation is unclear due to measurement constraints.
- Cuticle conductance to water vapor (*g*_*cw*_) was estimated from the difference between calculated and direct measurement of CO_2_ concentration in the leaf airspace (*C*_*i*_) of amphi-stomatous tobacco and sunflower. We estimated *g*_*cw*_ in a series of light and dark experiments, and partitioned *g*_*w,min*_ into cuticle and stomatal components. Some leaves were detached to simulate severe drought through desiccation conditions where *g*_*w,min*_ is generally determined.
- Between light and dark experiments each *g*_*cw*_ was in close agreement, and successfully corrected the discrepancies of calculations from direct measurements. In the dark, either stomatal or cuticle conductance dominated the *g*_*w,min*_, suggesting either of them can control the minimum water loss. In the detached leaves, *g*_*cw*_ could not be estimated likely due to unsaturation in the leaf airspace, and *g*_*w,min*_ was progressively underestimated.
- Besides cuticle, leaf water status is a potential pitfall of the standard gas exchange model. Our technique is useful to study the minimal gas exchange as well as to refine the model.

## Introduction

Land plants have to remain more or less turgid by balancing transpirational water loss with root water uptake. The epidermis and the cuticle covering the outer surface enables stomatal pores to actively control transpiration, but also limits CO_2_ uptake for photosynthesis. Therefore, stomata close to avoid both excessive water loss during daytime and dispensable water loss at nighttime. Nevertheless, water still escapes through the cuticle and imperfectly closed stomata. Such minimum transpiration can be considered part of plant water relations (Caird et al., 2007; Howard & Donovan, 2007; Resco de Dios et al., 2019). The quantification of minimal transpiration has recently gained a great deal of attention because of the broader impacts on water and carbon fluxes models in the scales from a single leaf to global vegetation (Hanson et al., 2016, Lombardozzi et al., 2017; Duursma et al., 2019. The diffusion pathway at the leaf-atmosphere interface could also have important implications for the deposition of air pollutants (Kerstiens & Lendzian, 1989; Musselman & Minnick, 2000; Barber et al., 2004), and emission of volatile organic compounds (Niinemets et al., 2004).

Minimum transpiration is controlled at the leaf by the minimum total conductance to water vapor (*g*_*w,min*_) composed of the cuticle conductance (*g*_*cw*_) and the minimum stomatal conductance to water vapor (*g*_*sw*_’_,*min*_) as:

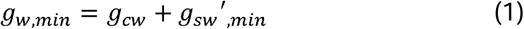

In general, two different methods are used to determine minimum conductance; measuring mass loss of detached leaves or measuring transpiration through gas exchange under conditions of assumed maximum stomatal closure (e.g., in darkness, application of abscisic acid, or under drought). Minimum conductance tends to be larger than the cuticle conductance measured in isolated astomatous cuticles/leaf surfaces, and the mass-loss method tends to have a smaller value than the gas-exchange method (Kersiens 1996; Duursma et al., 2019). These differences could be caused by leaky stomata or differences in vapor pressure difference between leaf airspace and atmosphere (VPD). The relative importance of leaky stomata is supported by a strong positive correlation between stomatal density and minimum conductance among sorghum genotypes (Muchow & Sinclair, 1989) and rice cultivars (Saito & Futakuchi, 2010). Also, a majority of water loss (50-95%) was observed from the stomatous side in hypostomatous conifers during desiccation (Brodribb et al., 2014).

However, control of minimum conductance is yet unclear due to constraints of these methods to assess cuticle components on the stomatous leaf surface. Santrucek et al. (2004) used mixtures of He, N_2_, and CO_2_ to alter water vapor diffusivity to segregate the water transport through the cuticle isolated from hypostomatous *Hedera helix* leaves. They found that the stomatous cuticle had 11 times larger conductance than the astomatous cuticle, arguing that thinner cuticles of the guard cells had the larger conductance. This can be an alternative explanation for the evidence of leaky stomata mentioned above. Moreover, Boyer (2015a) measured gas exchange on the astomatous surface along with leaf water status in grape leaves and found that *g*_*cw*_ declined as turgor decreased. This implies that the intact cuticles would be more conductive when turgor is high (e.g. during nighttime) than when turgor is low (e.g. during daytime and under drought). Therefore, leaf water status can affect cuticle conductance as well as stomatal leakiness. Overall, both *g*_*cw*_ and *g*_*sw*_’_,*min*_ can be variable, potentially accounting for the discrepancy among the measurement techniques and sample conditions.

In order to understand the controls of minimal transpiration, we must separate each component in intact leaves. In a companion work (MS#1), we developed a gas exchange technique to estimate *g*_*cw*_ on stomatous leaf surfaces. By applying the technique in tobacco and sunflower leaves, the present study attempted to partition the *g*_*w,min*_ into *g*_*cw*_ and *g*_*sw*_’_,*min*_. To validate the technique, experiments were performed with attached leaves in dark and were compared with experiments in light (MS#1). In addition, we desiccated leaves by cutting the petioles to imitate the mass-loss method while gas exchange was measured. The comparison of cuticle conductance between light and dark validated the method and showed the mass-loss method underestimated the minimum conductance, likely due to unsaturation of leaf intercellular airspace during leaf drying.

## Materials and methods

### Plant material

Tobacco (*N. tabacum* L. cv. Samsun) and sunflower (*H. annuus* L. cv. Mammoth Russian) plants were grown in a greenhouse located at the University of New Mexico [35.08°N, 106.62°W, 1587 m.a.s.l] under ambient CO_2_ at 24.2/19.9 °C day/night temperature as described in Tominaga et al. (MS#1). All experiments with these plants were conducted with upper fully expanded leaves.

### Dual gas exchange system

We used an open gas exchange system (LI-6800; LI-COR, Lincoln, NE, USA) combined with a closed system which measures intercellular CO_2_ concentration in leaves (*C*_*i*_) directly (for more details, see MS#1). Briefly, the LI-6800 system equipped with a large 6 x 6 cm chamber with a LED light source (6800-13L; LI-COR) measured gas exchange on the adaxial side while the infrared gas analyzer (IRGA; LI-7000; LI-COR) in the closed loop connected to the same leaf area of the opposite abaxial side measured CO_2_ concentration equilibrated with that inside the leaf (measured *C*_*i*_; *C*_*i(m)*_). The *C*_*i*_ was also calculated (*C*_*i(c)*_) from the gas exchange measurements according to a standard procedure (von Caemmerer & Farquhar, 1981). The advantage in this dual system is that conductance of CO_2_ can be determined independently of conductance of water vapor (Boyer & Kawamitsu, 2011). Because these independent measurements are simultaneous, discrepancy between the two conductances should be implicated in the different diffusion properties between the CO_2_ and water (Boyer & Kawamitsu, 2011; Boyer, 2015b; Tominaga & Kawamitsu 2015; Tominaga et al., 2018). We used this discrepancy to isolate the cuticle conductance (*g*_*cw*_) (see below).

### Cuticle conductance and minimum conductance

Using the combined gas-exchange system described above, the stomatal conductance to CO_2_ (*g*_*sc*_’) and that to water vapor (*g*_*sw*_) can be estimated from the measurements of CO_2_ flux (*A*) and water vapor flux (*E*), respectively as:

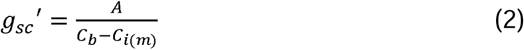

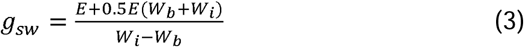

where the *C*_*b*_ and the *W*_*b*_ are CO_2_ and water vapor concentrations in the boundary layer on the adaxial leaf surface respectively, and the *W*_*i*_ is water vapor concentration inside the leaf, which is generally assumed saturated at the leaf temperature. In our companion work (MS#1)—light induction experiments, gas exchange was measured with the same equipment, plant materials, and calculations. When plotting *g*_*sw*_ against *g*_*sc*_’ during the initial response (<10 min), their regression line had a positive y-intercept. Because CO_2_ hardly moves across the cuticle and epidermis (Boyer et al., 1997; Boyer, 2015a), *g*_*sc*_’ would represent CO_2_ conductance entirely through stomata. Considering that *g*_*sc*_’ becomes zero when the stomata close completely, we extrapolated *g*_*cw*_ as the y-intercept. In the present experiments *g*_*cw*_ was similarly estimated from the regression line while stomata closed in dark or desiccation (while the *g*_*sc*_’ approached the minimum). In all the experiments, photosynthesis was fully induced under irradiance of either 200 and 1200 µmol m^-2^ s^-1^ photosynthetically active radiation (PAR) prior to the dark and 200-300 µmol m^-2^ s^-1^ PAR prior to desiccation, respectively.

Because *E* is composed of stomatal transpiration plus cuticular transpiration, the minimum *g*_*sw*_ derived from Eq. (3) was the minimum total conductance (*g*_*w,min*_) in this study. In turn, the actual minimum stomatal conductance (*g*_*sw*_’_,*min*_) was estimated by subtracting the *g*_*cw*_ from the *g*_*w,min*_, according to Eq.(1). A summary of parameters referred to within the text is shown with accompanying units in Table 1.

**Table 1.** A summary of parameters referred to within the text.

| Parameter | Definition | Units |
| --- | --- | --- |
| $A$ | Net CO<sub>2</sub> assimilation rate | $\mu\text{mol m}^{-2} \text{s}^{-1}$ |
| $C_b$ | Boundary layer CO<sub>2</sub> concentration | $\mu\text{mol mol}^{-1}$ |
| $C_i$ | Intercellular CO<sub>2</sub> concentration | $\mu\text{mol mol}^{-1}$ |
| $C_{i(c)}$ | Calculated $C_i$ | $\mu\text{mol mol}^{-1}$ |
| $C_{i(m)}$ | Directly measured $C_i$ | $\mu\text{mol mol}^{-1}$ |
| $C_{i(c),cut}$ | $C_{i(c)}$ corrected for $g_{cw}$ | $\mu\text{mol mol}^{-1}$ |
| $\Delta C_i$ | Difference between $C_{i(c),cut}$ and $C_{i(m)}$ ($C_{i(m)} - C_{i(c),cut}$) | $\mu\text{mol mol}^{-1}$ |
| $E$ | Transpiration rate ($E_c + E_s$) | $\text{mmol m}^{-2} \text{s}^{-1}$ |
| $E_c$ | Cuticular transpiration rate | $\text{mmol m}^{-2} \text{s}^{-1}$ |
| $E_s$ | Stomatal transpiration rate | $\text{mmol m}^{-2} \text{s}^{-1}$ |
| $g_{cw}$ | Cuticle conductance to water vapor | $\text{mmol m}^{-2} \text{s}^{-1}$ |
| $g_{sc}'$ | Stomatal conductance to CO<sub>2</sub> estimated from $C_{i(m)}$ | $\text{mmol m}^{-2} \text{s}^{-1}$ |
| $g_{sw}$ | Stomatal and cuticular conductance to water vapor | $\text{mmol m}^{-2} \text{s}^{-1}$ |
| $g_{sw}'$ | Stomatal conductance to water vapor | $\text{mmol m}^{-2} \text{s}^{-1}$ |
| $g_{sw}',min$ | Minimum stomatal conductance to water vapor | $\text{mmol m}^{-2} \text{s}^{-1}$ |
| $g_{w,min}$ | Minimum total conductance to water vapor ($g_{sw}',min + g_{cw}$) | $\text{mmol m}^{-2} \text{s}^{-1}$ |
| PAR | Photosynthetically active radiation | $\mu\text{mol m}^{-2} \text{s}^{-1}$ |
| RH | Relative humidity | % |
| VPD | Vapor pressure difference between leaf airspace and atmosphere | kPa |
| $W_b$ | Boundary layer water vapor concentration | $\text{mmol mol}^{-1}$ |
| $W_i$ | Intercellular water vapor concentration calculated at leaf temperature with assuming saturation | $\text{mmol mol}^{-1}$ |
| $W_i'$ | Intercellular water vapor concentration calculated from $\Delta C_i$ with assuming unsaturation | $\text{mmol mol}^{-1}$ |

### Intercellular CO_2_ gradients

Vertical gradients of CO_2_ were estimated from the anatomically determined intercellular CO_2_ conductance as described in Tominaga et al. (MS#1).

### Statistics

The number of replications is presented in figure legends for each experiment. The results are given as means with SDs.

## Results

### Dark

After photosynthesis was fully induced from dark, the light was turned off and the direction of CO_2_ flux or net assimilation rate (*A*) was reversed from positive to negative (Fig. 1a). Stomatal conductance to CO_2_ (*g*_*sc*_) and water vapor (*g*_*sw*_) changed in parallel, reflecting the stomatal dynamics (Fig. 1b). Compared to photosynthesis, the stomatal response was slower. The closure in the dark was relatively faster in the first 10-30 min, and then became gradual (Fig. 1b). Calculated *C*_*i*_ (*C*_*i(c)*_) was larger than measured *C*_*i*_ (*C*_*i(m)*_) during the early induction, and the difference decreased as stomata opened (insets of Fig. 1c). In the subsequent darkness, the *C*_*i(m)*_ departed from the *C*_*i(c)*_ as the stomata closed while the *C*_*i(c)*_ remained relatively constant (Fig. 1c).

**Fig. 1.**
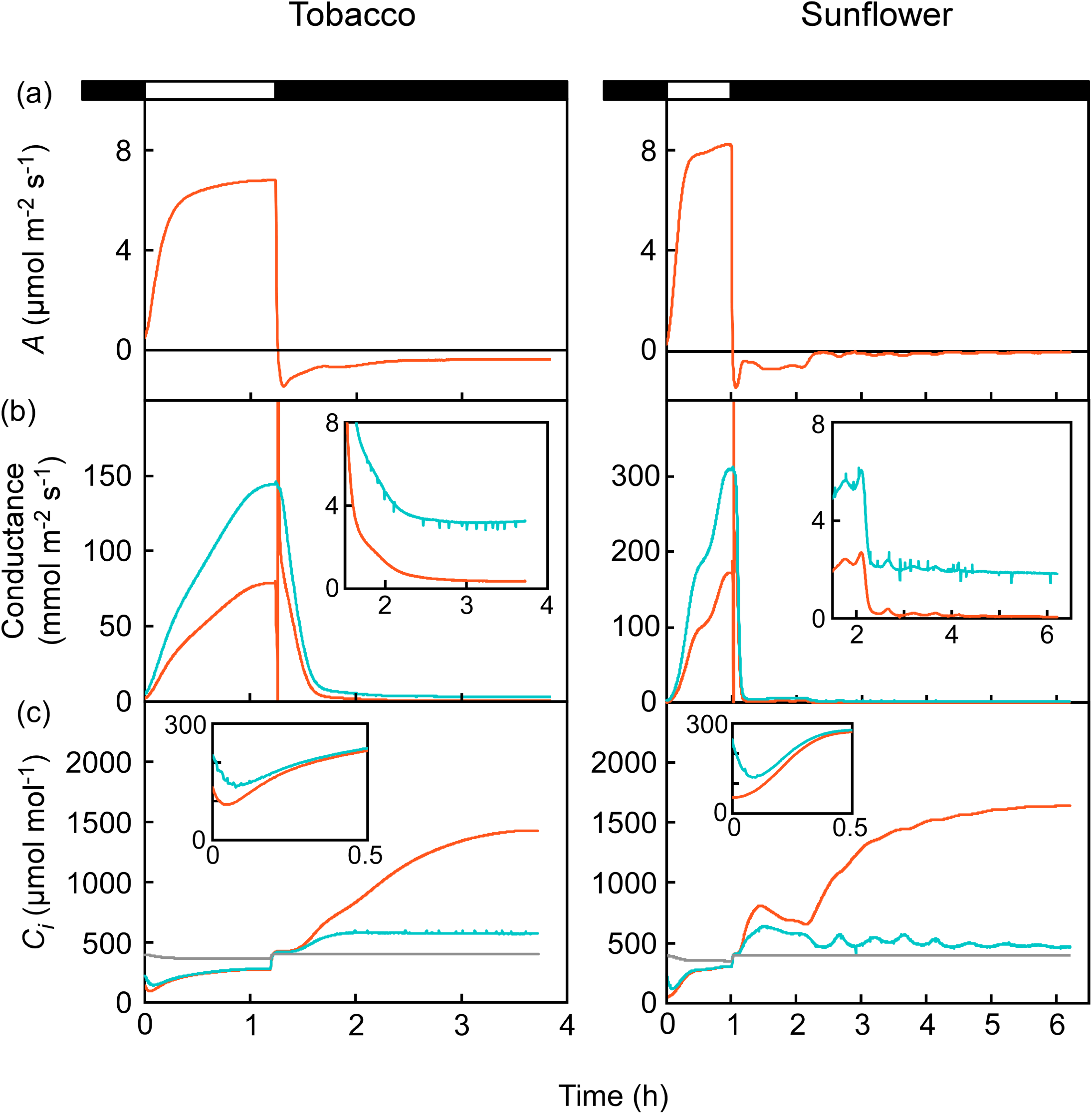
Changes in the gas exchange parameters during the light and dark experiments. Assimilation rate (*A*) (a), stomatal conductances of water vapor (*g*_*sw*_, blue line) and CO_2_ (*g*_*sc*_’, orange line) (b), and calculated intercellular CO_2_ concentration (*C*_*i(c)*_, blue line), measured intercellular CO_2_ concentration (*C*_*i(m)*_, orange line), and ambient CO_2_ concentration (*C*_*a*_, grey line) (c) after dark-adapted leaf of tobacco (left) and sunflower (right) were illuminated with 200 µmol m^−2^ s^−1^ PAR and then re-darkened (indicated as white and black columns on top of (a), respectively). The *g*_*sw*_ was derived from the standard calculation based on the water vapor flux (Eq. (2)), whereas the *g*_*sc*_’ was derived from the direct measurement based on the CO_2_ flux (Eq. (3)). In (b), data when the values approached to the minimum in the dark was zoomed in the inset. In (c), data during 0.5 h after the light induction was zoomed in the inset. Note both *C*_*i(c)*_ *and C*_*i(m)*_ were above *C*_*a*_ in dark. Representative experiments from ten to eight replications in tobacco and sunflower, respectively.

During the initial induction, a linear relationship was found between *g*_*sw*_ and *g*_*sc*_’ (Fig. 2). Similarly, the linear relationship was observed in the subsequent dark while the *g*_*sw*_ was approaching the minimum (Fig. 2). Assuming that stomata close completely when *g*_*sc*_’ is zero, we extrapolated the *g*_*cw*_ as the y-intercept of the regression. Correction for these *g*_*cw*_ estimates moved the *C*_*i(c)*_ closer to the *C*_*i(m)*_ in the dark (*C*_*i(c),cut*_ in Fig. 3) as can be seen in the light (insets of Fig. 3). In these leaves, the intercellular CO_2_ gradient did not account for the difference between the *C*_*i(c)*_ and *C*_*i(m)*_ during the early induction (MS#1). Likewise, the gradients were negligible in the dark (Fig. S1). These results further supported the view that cuticle conductance to water vapor is the major source of error in calculating *C*_*i*_ when stomatal conductance is small (MS#1). Since calculated *g*_*sw*_ includes *g*_*cw*_, it overestimates CO_2_ movement through the stomata. As a result, *C*_*i*_ is either overestimated when the CO_2_ diffuses into the leaf or underestimated when the CO_2_ diffuses out of the leaf.

**Fig. 2.**
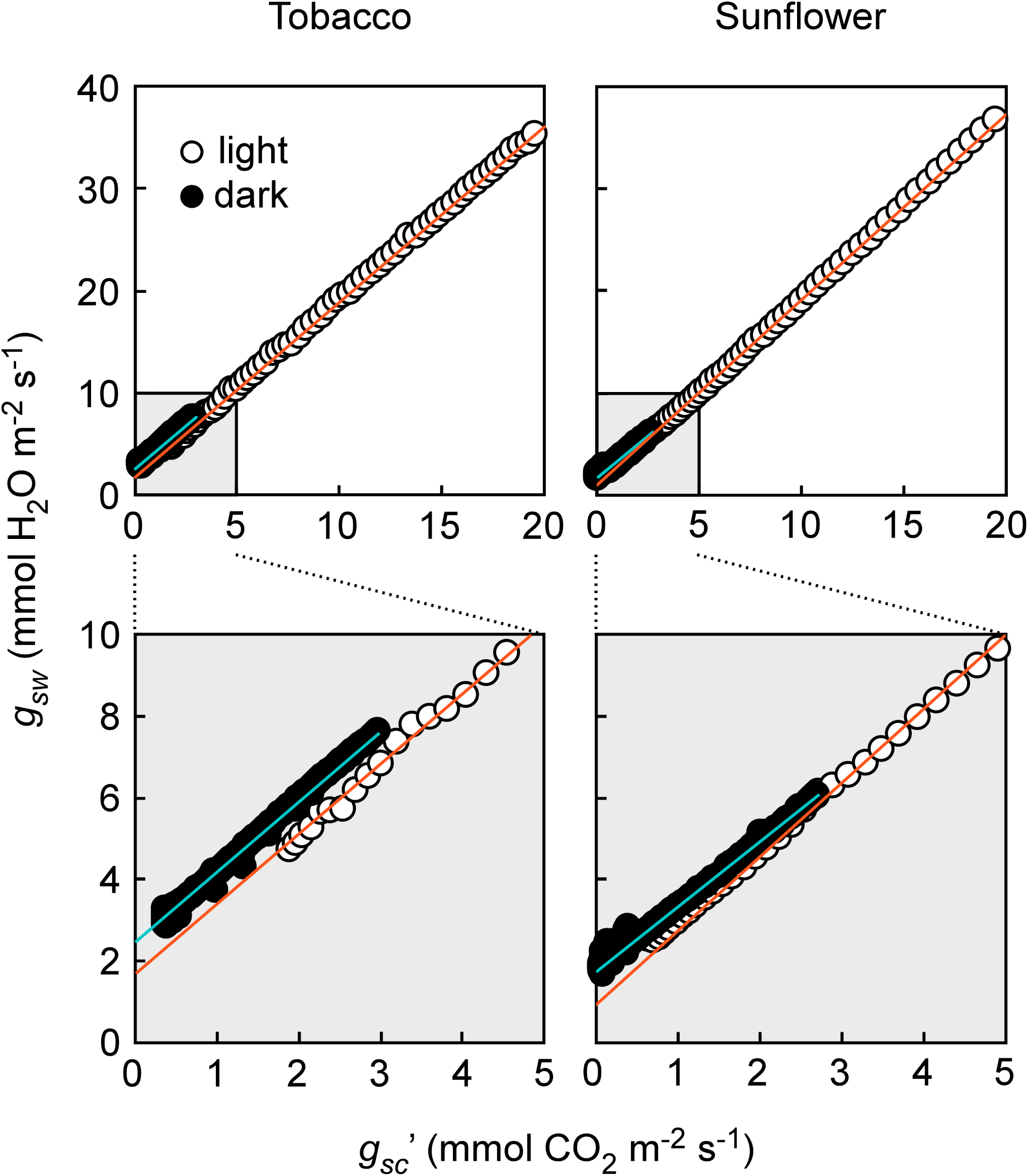
Relationships between *g*_*sw*_ and *g*_*sc*_’ during initial phase (*g*_*sc*_’ < 20 mmol CO_2_ m^−2^ s^−1^) of the light induction from the dark (open circle) and during the subsequent dark (closed circle) where *g*_*sc*_’ was approaching to the minimum (*g*_*sc*_’ < 3 mmol CO_2_ m^−2^ s^−1^). The Data for Fig. 1 is shown. The smaller range of stomatal conductances shown with the grey background (upper) are zoomed (lower). Shown regressions (orange and blue lines) are *Y*=1.71*X*+1.71 (*R*^*2*^=0.9997) for light and *Y*=1.70*X*+2.50 (*R*^*2*^=0.9944) for dark in tobacco, and *Y*=1.59*X*+1.75 (*R*^*2*^=0.9939) for light and *Y*=1.81*X*+0.94 (*R*^*2*^=0.9997) for dark in sunflower.

**Fig. 3.**
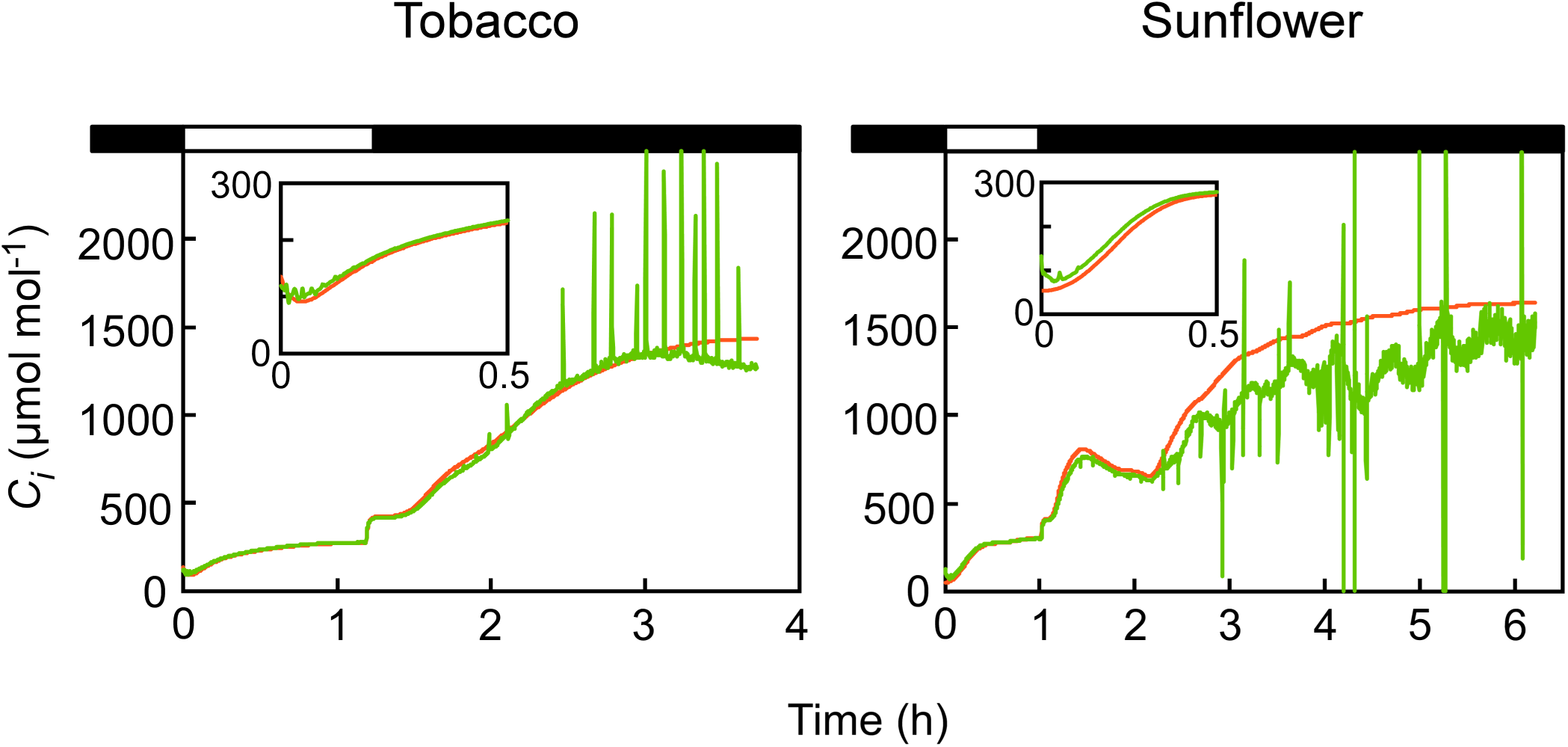
Correction for cuticle conductance to water vapor. Data for Fig. 1 is shown. The *C*_*i(c)*_ was corrected with the *g*_*cw*_ (*C*_*i(c),cut*_, green line) estimated for the light induction and for the subsequent dark, respectively, as shown in Fig. 2.

Despite a large variation of *g*_*cw*_ among leaves in both species, each leaf showed a good agreement in the *g*_*cw*_ between dark and light with *g*_*cw*_ values slightly but consistently larger in the dark (Fig. 4a), probably mediated through a change in leaf turgor (Boyer et al., 1997; Boyer, 2015a). In addition, the slope of the regression tended to be smaller in the dark and larger in the light in sunflower leaves (Fig. 4b). Those leaves had slope values close to 1.6 in the dark, matching the modeled gas diffusivity ratio of CO_2_ and water vapor for stomata (von Caemmerer & Farquhar, 1981: Massman, 1998). In the light, non-uniform stomatal openings might have affected the slope in sunflower (MS#1) though the potential effect was marginal (residual over-estimation in insets of Figs. 3 & S1).

**Fig. 4.**
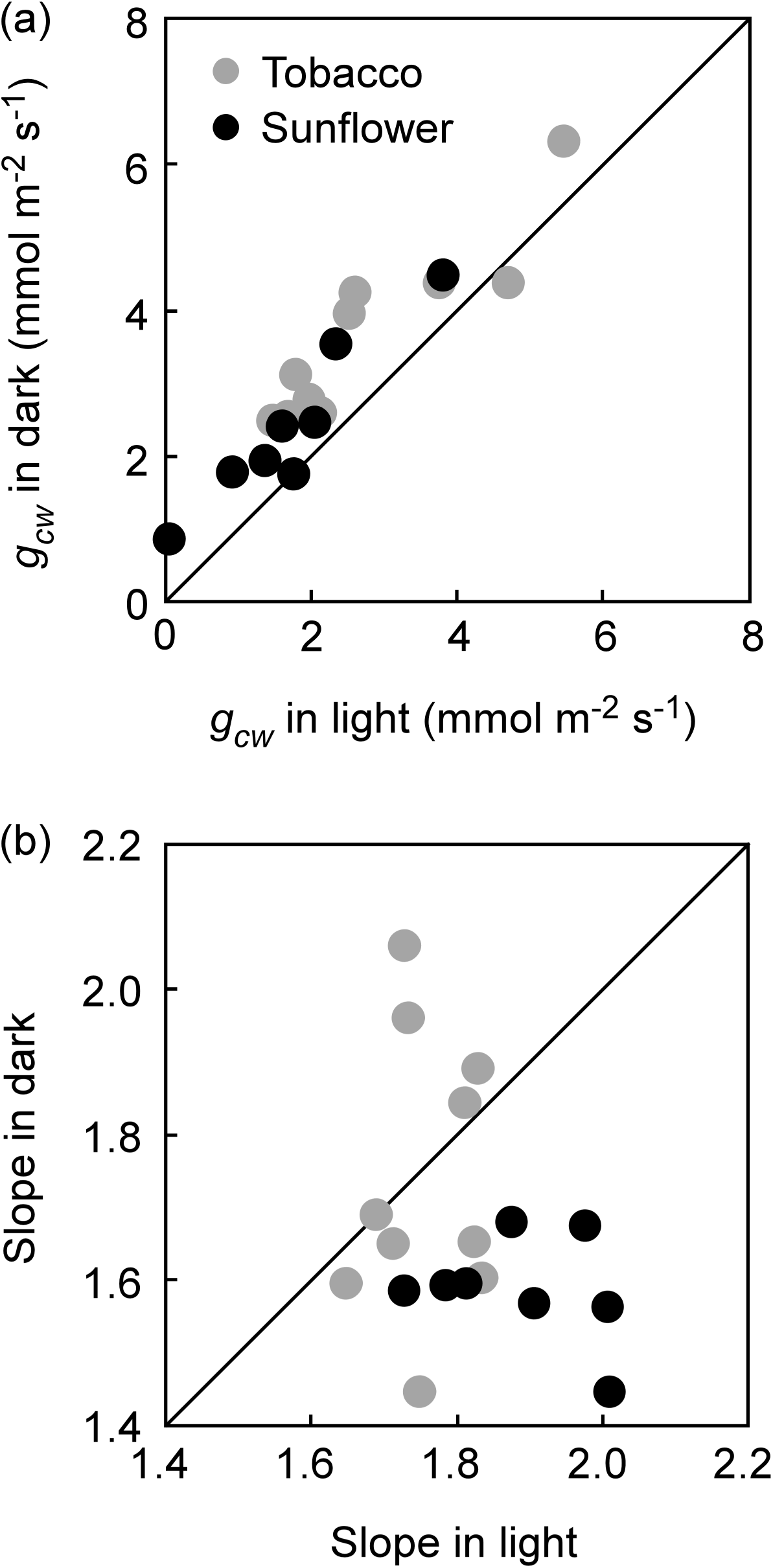
Relationships in *g*_*cw*_ (a) and in regression slope (b) between dark and light for tobacco (open circle) and sunflower (closed circle). Data above the 1:1 line indicates the larger values in dark than in light whereas data below the 1:1 line indicates the lower value in dark than in light.

We then calculated the minimum stomatal conductance (*g*_*sw*_’_,*min*_) by subtracting the *g*_*cw*_ from the total conductance to water in the dark (*g*_*w,min*_), according to Eq. (1). A positive correlation between the *g*_*w,min*_ and the *g*_*sw*_’_,*min*_ demonstrated that leaky stomata contributed to the minimum water loss (Fig. 5). At a severe stomatal closure, cuticle conductance could also have a large impact on the minimum transpiration. For example, when the *g*_*sw*_’_,*min*_ was less than 3 mmol m^-2^ s^-1^, the *g*_*cw*_ accounted as much as 77 ± 16% and 71 ± 32% of the *g*_*w,min*_ in tobacco and sunflower, respectively.

**Fig. 5.**
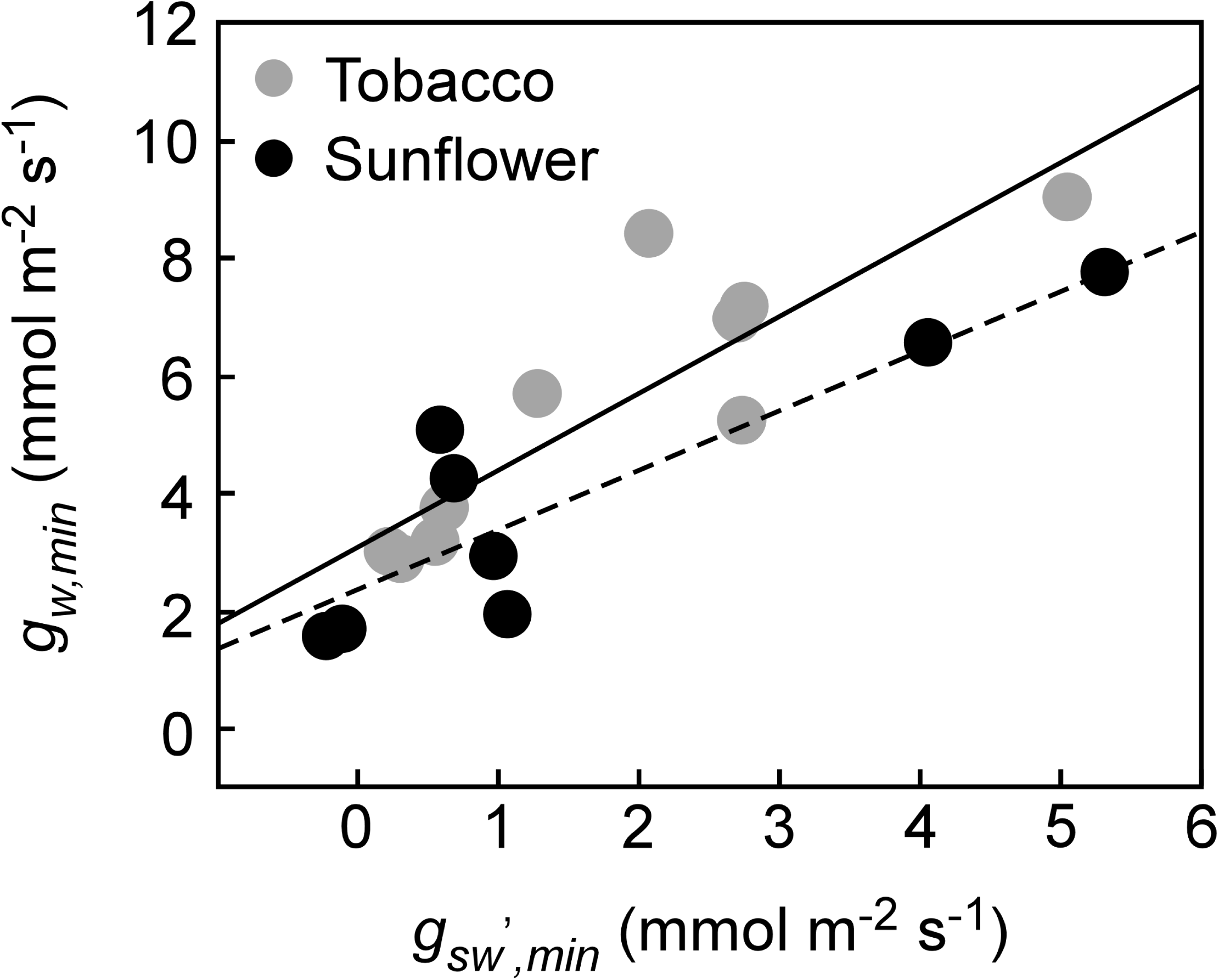
Relationships between *g*_*w,min*_ and *g*_*sw*_’_,*min*_ in dark. Solid and broken lines indicate regression lines for tobacco (*Y*=1.30*X*+3.10, *R*^*2*^=0.760, *n*=10) and sunflower (*Y*=1.01*X*+2.36, *R*^*2*^=0.766, *n*=8), respectively.

### Desiccation

We simulated a condition where minimum conductance is determined from mass loss of detached leaves while measuring the gas exchange in light (Fig. 6). Similar to the dark, stomata closed after a transitory opening immediately after the leaf was detached (Fig. 6b), then the leaf gradually lost water, eventually passing the turgor loss point. Positive *A* indicated that the CO_2_ diffused into the leaves (Fig. 6a), and so *C*_*i(c)*_ was expected to be overestimated by the cuticle conductance. However, the *C*_*i(c)*_ became lower than the *C*_*i(m)*_ as stomata closed while desiccation continued (Fig. 6c). When the *C*_*i(c)*_ was corrected by the *g*_*cw*_ estimated in the light for the same leaves (*C*_*i(c),cut*_), it further departed from the *C*_*i(m)*_ (Fig. 6c). The lower *C*_*i(c)*_ clearly suggested a significant source of error existed beyond cuticular conductance. As opposed to the cuticle effect, the *C*_*i(c)*_ underestimates actual *C*_*i*_ when the calculated *g*_*sw*_ is lower than the actual *g*_*sw*_.

**Fig. 6.**
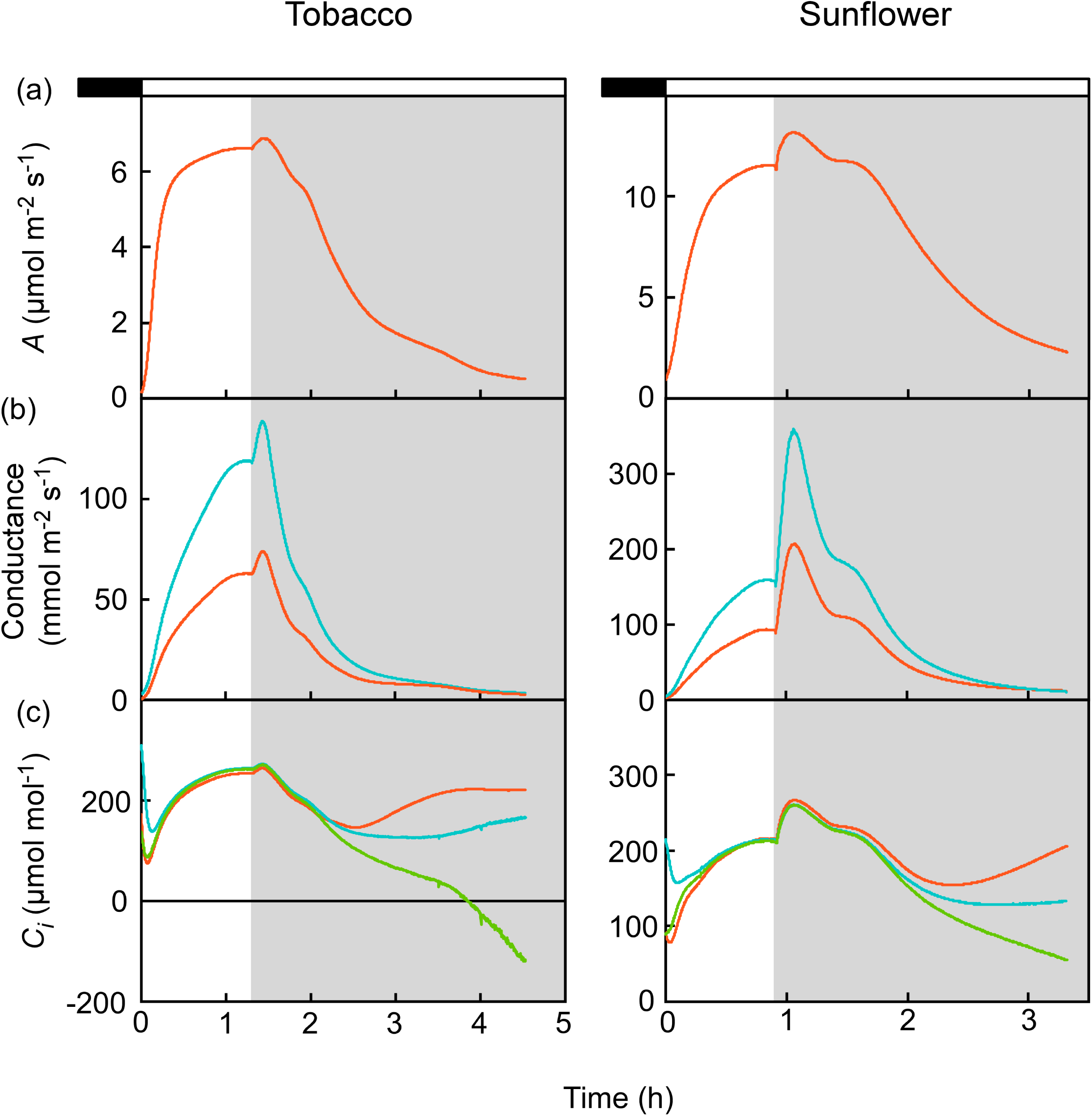
Change in the gas exchange parameters after dark-adapted leaves were illuminated with 200-300 µmol m^−2^ s^−1^ PAR and then petioles were cut (grey background). The *C*_*i(c)*_ was corrected only with the *g*_*cw*_ estimated for the light induction (*C*_*i(c),cut*_). Representative experiments from four replications in both species.

Since water re-supply to evaporating surfaces was limited as a result of detaching the leaf, a likely candidate causing the under-estimation of *g*_*sw*_ would be the unsaturation of water vapor in the intercellular airspace (*W*_*i*_). The calculation of *W*_*i*_ always assumes saturation or 100% relative humidity (RH). Using this assumption, actual RH could be estimated from the discrepancy between the calculation and the direct measurement of *C*_*i*_, as shown in Eq (A8) in the Appendix, and then converted to water potential (ψ) of the evaporating surfaces according to Eq. (A9). Plotting *C*_*i(c),cut*_ vs *C*_*i(m)*_, we found that *C*_*i(c),cut*_ started to divert ∼30 min after cutting the petiole (Fig. S2). It was then considered that the RH began to decrease (green lines in Fig. 7ab). This lagged behind the decrease of stomatal conductance beginning ∼10 min after the leaf was excised (Fig. 6b, blue lines in Fig. 7). The ψ could be as low as *–*62 MPa and *–*26 MPa in detached tobacco and sunflower leaves, respectively (Fig. 7b), as they continued to dry out but still maintained positive net *A. A* vs *C*_*i(m)*_ relationships indicated that photosynthesis started to be impaired ∼10-20 min after decreasing the ψ (Fig. S3, red lines in Fig. 7). Eventually, CO_2_ demand for photosynthesis fell short of CO_2_ supply, increasing the *C*_*i(m)*_ ∼50-60 min after the ψ began to decrease (Fig. 6c, purple lines in Fig. 7). When the *C*_*i(m)*_ started to increase the ψ reached *–*11.7 and *–*7.7 MPa in detached tobacco and sunflower leaves, respectively (Figs. 7b).

**Fig. 7.**
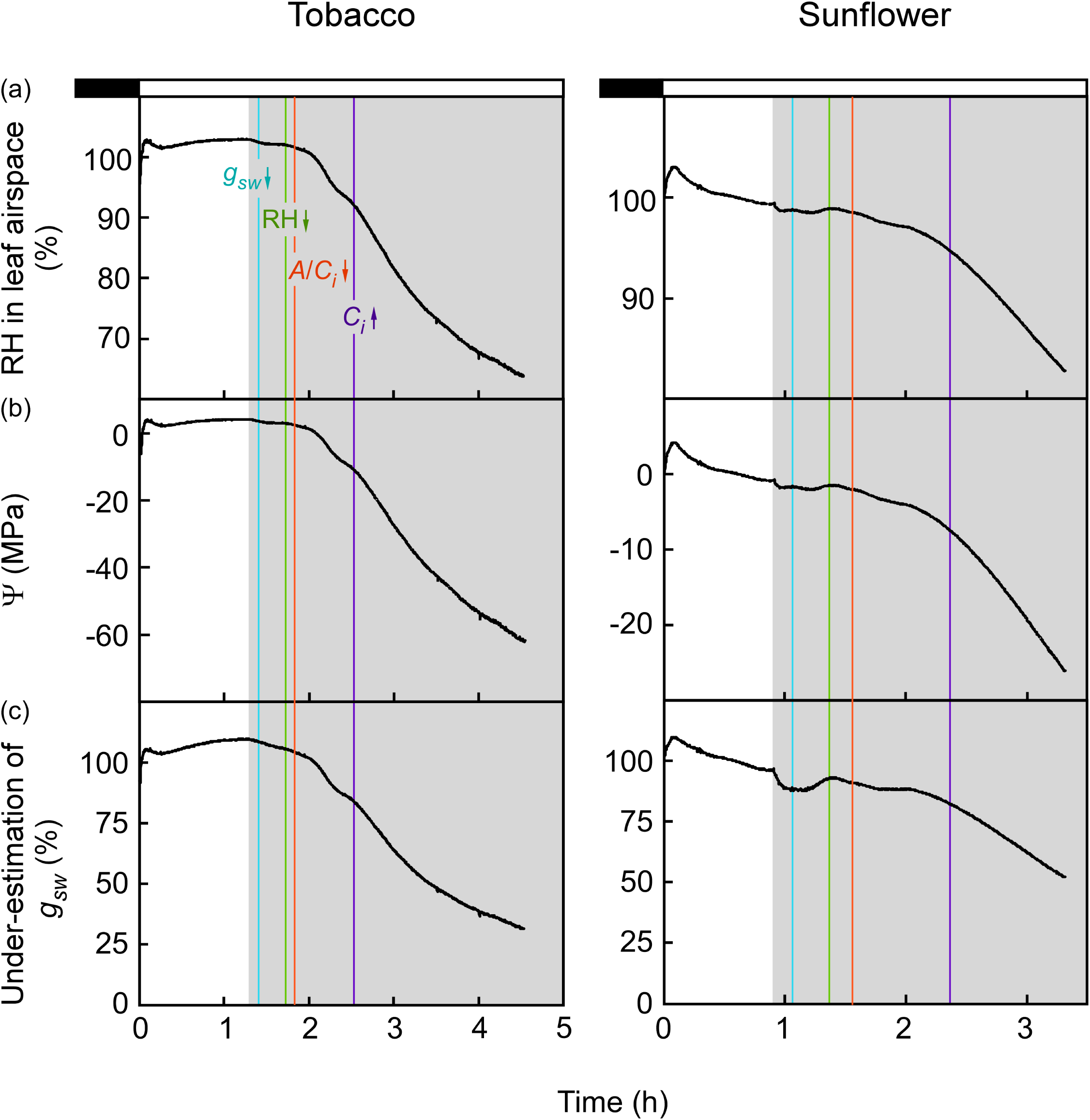
Change in relative humidity (RH) in leaf airspace (a), water potentials (Ψ) of evaporating surface (b), and under-estimation of *g*_*sw*_ (c). Data for Fig. 6 is shown. The RH was estimated according to Eq. (A6) whereas the Ψ was estimated from the RH according to Eq. (A8). The under-estimation of *g*_*sw*_ (%) was calculated as the proportion of saturation-based *g*_*sw*_ to unsaturation-based *g*_*sw*_. Different color lines indicate time-stamps for the initiation of physiological response to the leaf detachment: blue for the stomatal closure (Fig. 6b), green for the unsaturation (Fig. S2), orange for the suppression of photosynthesis (Fig. S3), and purple for the inflection of *C*_*i*_ (Fig. 6c).

Finally, the *g*_*sw*_ under 100% RH (i.e., standard calculation) was compared to the *g*_*sw*_ under the sub-saturated RH (Fig. 7c). As expected, the *g*_*sw*_ was progressively underestimated with decreasing the RH. Consequently, the *g*_*w,min*_ would be underestimated to 32% and 53% of the actual *g*_*w,min*_ in tobacco and sunflower, respectively (Fig. 7c).

## Discussion

Using the dual gas-exchange system, we estimated cuticle conductance on the adaxial stomatous side of intact leaves across a range of small stomatal conductances. This approach reduces uncertainties for determining cuticle conductance and can be achieved by measuring during the initial phase of light induction where the stomata that were initially closed in the dark gradually open in the light (MS#1). Here, we employed the technique to the transition from light to dark while stomata were closing. In contrast to light induction, calculations underestimate actual *C*_*i*_ in the dark as cuticle conductance to water causes an overestimation of *g*_*sw*_ and the ease that CO_2_ can diffuse out of the leaf (Fig. 1). The correction for the cuticle effect (Fig. 3) and the strong correlation of *g*_*cw*_ between light and dark (Fig. 4) together demonstrate the robustness of our approach. By comparing *g*_*sw*_ and *g*_*sc*_’ across a range of low conductances, we were able to make small extrapolations to the condition of complete stomatal closure for calculating *g*_*cw*_, thereby solving the challenge presented by leaky stomata. Although the *g*_*cw*_ values scattered by several-fold within the similar range in both tobacco and sunflower (MS#1), isolated *g*_*cw*_ has been shown to vary among species by over three orders of magnitude (Kerstiens, 1996; Kerstiens 2006; Schuster et al., 2017). Our future work will test if such a broad range of *g*_*cw*_ can also be seen in intact cuticle.

Estimation of *g*_*cw*_ on the stomatous leaf surface allowed partitioning of minimum conductance into cuticular and stomatal components in the dark. According to Eq. (1), *g*_*w,min*_ is more controlled by stomatal than cuticular water vapor loss when *g*_*sw*_’_,*min*_ is larger than *g*_*cw*_ and vice versa. In our experiments *g*_*sw*_’_,*min*_ was found to be both larger or smaller than the *g*_*cw*_ depending on the conditions, suggesting that both cuticle and leaky stomata are important in controlling the minimal transpiration. In some leaves the *g*_*sw*_’_,*min*_ could be so small that *g*_*cw*_ accounted more than 80% of the *g*_*w,min*_. In addition, the *g*_*sc*_’ could be so small that respired CO_2_ was accumulated inside the leaf as much as several-fold higher than the atmospheric [CO_2_] of 400 ppm (Fig. 1). Similar results were also obtained in previous direct *C*_*i*_ measurements during nighttime in sunflower leaves (Boyer & Kawamitsu, 2011).

While the strong correlation of the *g*_*cw*_ between dark and light suggests a stable property of the solid cuticle, a slight increase of the *g*_*cw*_ in the dark was also evident (Fig. 4a). We speculate that the leaves became more turgid when transpiration was low in the dark, and the cuticle stretched. The stretching would then increase the conductance by shortening the diffusion path-length of water molecules in the waxes (Boyer, 2015a). However, this will require direct tests from future studies.

### Underestimation of minimum conductance due to unsaturation of leaf airspaces

In addition to nighttime measurements, minimum conductance can also be important for daytime measurements under drought conditions that are severe enough to limit re-supply to leaves. We simulated this by detaching photosynthetically active illuminated leaves and monitoring conductance and assimilation through the initial rapid increase caused by the release of tension, and subsequent rapid and gradual decreases as stomata closed and leaves dried. Surprisingly, we were unable to estimate the cuticle conductance accurately soon after stomata began closing and *C*_*i(c)*_ became lower than *C*_*i(m)*_, even though assimilation was near 50% of maximum values (Fig. 6). This contrasts with the leaves fed with ABA in which *C*_*i(c)*_ was increasingly higher than *C*_*i(m)*_ as stomata closed (Boyer, 2015b, Tominaga & Kawamitsu 2015). These opposite patterns suggest that conditions limiting hydraulic conductance to leaves can cause further difficulties for calculating *C*_*i*_. This may have implications for how conductance and assimilation should be modeled during drought stress, potentially providing a new avenue for accurately integrating hydraulic conductance.

We found that the under-estimation by *C*_*i(c)*_ can be explained by allowing the calculated *W*_*i*_ to become progressively sub-saturated as the detached leaf loses water that cannot be re-supplied through the vasculature (Figs. 7 and S4). Importantly, if the assumption that *W*_*i*_ is saturated during drying is used, then minimum conductance will be underestimated. Therefore, it is critical to know the dynamics of *W*_*i*_ saturation during drying for both the mass-loss method and the gas exchange method since the methods share assumptions about *W*_*i*_ saturation (Burghardt & Riederer, 2006). More often than not, *g*_*w,min*_ determined from the mass-loss method is regarded as *g*_*cw*_ (Schuster et al., 2017), which is problematic for understanding what controls water loss when stomata are mostly closed. Notably, unsaturation should have a smaller effect on *g*_*cw*_ calculations than on the *g*_*w,min*_ in the mass-loss method. This is because the *W*_*i*_ used for *g*_*cw*_ represents the surface conditions below the cuticle, whereas the *W*_*i*_ used for *g*_*w,min*_ represents the sub-stomatal cavity (Kerstiens, 1996). When we assumed that the *W*_*i*_ used for *g*_*cw*_ remains saturated (Eq. A4) during our drying experiment, but allowed *W*_*i*_ in the sub-stomatal cavity to change, it effectively corrected the calculation of *C*_*i*_ (Fig. S4). Furthermore, when the stomata remain partially open as was seen in Fig. 6b, the stomatal transpiration diminishes with decreasing VPD as *W*_*i*_ in the sub-stomatal cavity diminishes. While this decrease of *W*_*i*_ decreases the *g*_*sw*_’_,*min*_ and so the *g*_*w,min*_, the *g*_*w,min*_ increasingly approximates the *g*_*cw*_.

Our estimate of unsaturation of leaf airspaces during drying suggests that the water potential of leaf cell walls (i.e., apoplastic water) can be far more negative than is currently thought to be possible (Buckley and Sack, 2019). Indeed, our results estimated that apoplastic ψ decreased as low as *–*60 MPa while net photosynthesis was still positive, and potentially lower if the measurements continued (Fig. S5). This has significant implications for hydraulic conductance pathways inside the leaf. Our data also show a progression of response times to the leaf detachment in the order of *g*_*sw*_ *<* ψ < *C*_*i*_, which may indicate that the leaves initiated the stomatal closure to maintain the cellular homeostasis (Figs. 7b). Similar to our results, Cernusak et al. (2018) estimated that the relative humidity could be as low as 77% and 87% (ψ ≂ *–*35 MPa and *–*18 MPa, respectively) in *Pinus edulis* and *Juniperus monosperma* even when the leaves opened stomata and actively photosynthesized. Even if transient cavitation occurred in *P. edulis* and *J. monosperma*, enough to impair re-supply, there must be a mechanism to maintain the symplastic ψ—the cellular metabolism—while the water ‘film’ covering the cells had very low ψ. When and how such low apoplastic ψ occur have yet to be investigated. It is also unknown whether apoplastic ψ affects the cell turgor and/or cuticle conductance.

Since we have severed the leaf, it is conceivable that xylem water is supplied to the connected apoplast faster than the symplast since the xylem tension has been released. The increased apoplast water would be then re-distributed to the lower water potential (higher solute) symplast throughout the leaf. The initial effect of cutting on water movement can be seen with the transient increase in *g*_*sw*_ as water moves into the guard cells (Fig. 6b). Without the ability to re-supply water through xylem, negative water potential would be developed in apoplast with decreasing the matric potential as the water film gets thinner in the cell wall pores [with a dimeter of 4-6.5 nm (Kramer & Boyer, 1995)]. This might allow the formation of unexpected micro-scale water potential gradients in leaves, with the decline in photosynthesis illustrating the progressive drying of mesophyll cells into the interior of the leaf.

Until these effects can be easily corrected, the calculations must be viewed with skepticism. The dual gas-exchange system presented here enables a robust measurement of *C*_*i*_, and further provides an opportunity to study cuticle conductance and unsaturation, thereby refining general leaf gas-exchange models.

## Acknowledgements

We are deeply grateful to Prof. John S. Boyer (University of Missouri). This work was supported by funding to JT through the JSPS KAKENHI 18J00308 at Hiroshima University, and to DTH through the NSF EPSCoR Program under Award # IIA-1301346 and through NSF IOS 1658951 at the University of New Mexico. Any opinions, findings, and conclusions or recommendations expressed in this material are those of the authors and do not necessarily reflect the views of the National Science Foundation. JT is supported by Research Fellowships for Young Scientists from the Japanese Society for the Promotion of Science.s

## Author contributions

JT planned and designed the research. JT and JRS performed experiments and analyzed data. JT and DTH wrote the manuscript.

## Appendix

### Estimation of the intercellular humidity and water potentials in leaves

In the simplest form, *C*_*i(c)*_ would be expressed as (Moss & Rawlins, 1963):

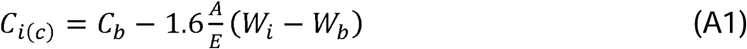

When correcting for cuticle effect, Eq. (A1) would become:

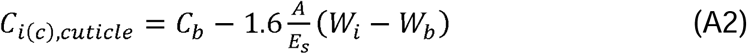

where *E*_*s*_ is the stomatal transpiration recalculated by subtracting the cuticular transpiration (*E*_*c*_) from the bulk leaf transpiration (*E*) as:

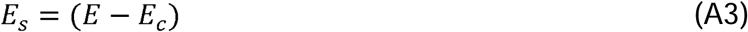

where *E*_*c*_ was estimated from the cuticle conductance as:

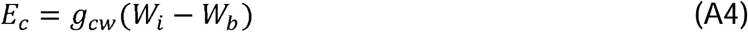

The *W*_*i*_ for cuticular transpiration in Eq. (A4) is assumed to be in between cuticular membrane and the epidermal cell wall (Kerstiens, 1996) whereas the *W*_*i*_ is assumed to be at the intercellular airspace for stomatal transpiration in Eqs. (3, A1, & A2). In practice, both are calculated from the same leaf temperature measurement, and thus are not distinguished. We assumed that the unsaturation altered stomatal *W*_*i*_ by affecting the vapor pressure of water in the air phase of the sub-stomatal but not the cuticular *W*_*i*_, which is essentially in liquid phase (Kerstiens, 1996), thus we used saturated *W*_*i*_ for calculating *E*_*c*_.

Similar to the Eq. (A2), *C*_*i(m)*_ would be expressed as:

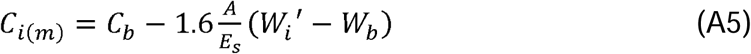

where *W*_*i*_’ is the actual water vapor concentration inside the leaf. Then, the difference between the *C*_*i(m)*_ and *C*_*i(c),cut*_ (Δ*C*_*i*_) is:

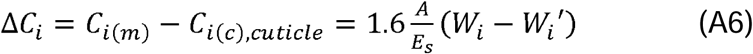

Eq. (A6) shows that under-estimation or unsaturation of *W*_*i*_ (*W*_*i*_ *– W*_*i*_’) increases as under-estimation of *C*_*i*_ (*C*_*i(m)*_ *– C*_*i(c),cut*_) increases. Rearranging Eq. (A6) gives:

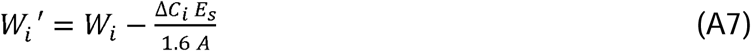

Relative humidity (RH) in the intercellular airspace is then calculated as:

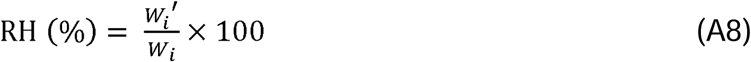

Although Eqs. (A1 & A2) can be expressed in a more comprehensive version for the standard calculations elaborated by von Caemmerer and Farquhar (1981), the Eq. (A6) corrected the *C*_*i(c)*_ derived from the standard calculations almost completely (Fig. S4), suggesting that the model difference hardly affected the estimations of RH shown in figure 7a.

At equilibrium, the water potential (ψ) of an aqueous solution is a function of the RH over the solution as (Kramer & Boyer, 1995):

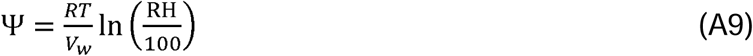

where *V*_*w*_ is the molar volume of water (1.8·10^−5^ mol m^−3^), *R* is the gas constant (8.314·10^−6^ MPa m^3^ mol^−1^ K^−1^) and *T* is temperature in kelvins.

In leaves showing positive net *A*, under-estimation of *W*_*i*_ is indicated when Δ*C*_*i*_ is positive, according to Eq. (A6). When Δ*C*_*i*_ is negative, on the other hand, *W*_*i*_ ‘will be above saturation (*W*_*i*_ ‘> *W*_*i*_), and ψ will be greater than 0. Although these conditions are impossible, it happened because the calculated *C*_*i*_ was still larger than the *C*_*i(m)*_ after correcting the *g*_*cw*_ (*C*_*i(c),cut*_ > *C*_*i(m)*_) (Figs. 6 & 7). It was likely that under-estimation of *W*_*i*_

was masked by factors other than cuticle, such as vertical *C*_*i*_ gradients and non-uniform stomatal apertures (MS#1). Despite these adverse effects, under-estimation of *W*_*i*_ was evident.

