## Supplementary material for "Minimum conductance in leaves—cuticle, leaky stomata, or water vapor saturation?": Fig. S1

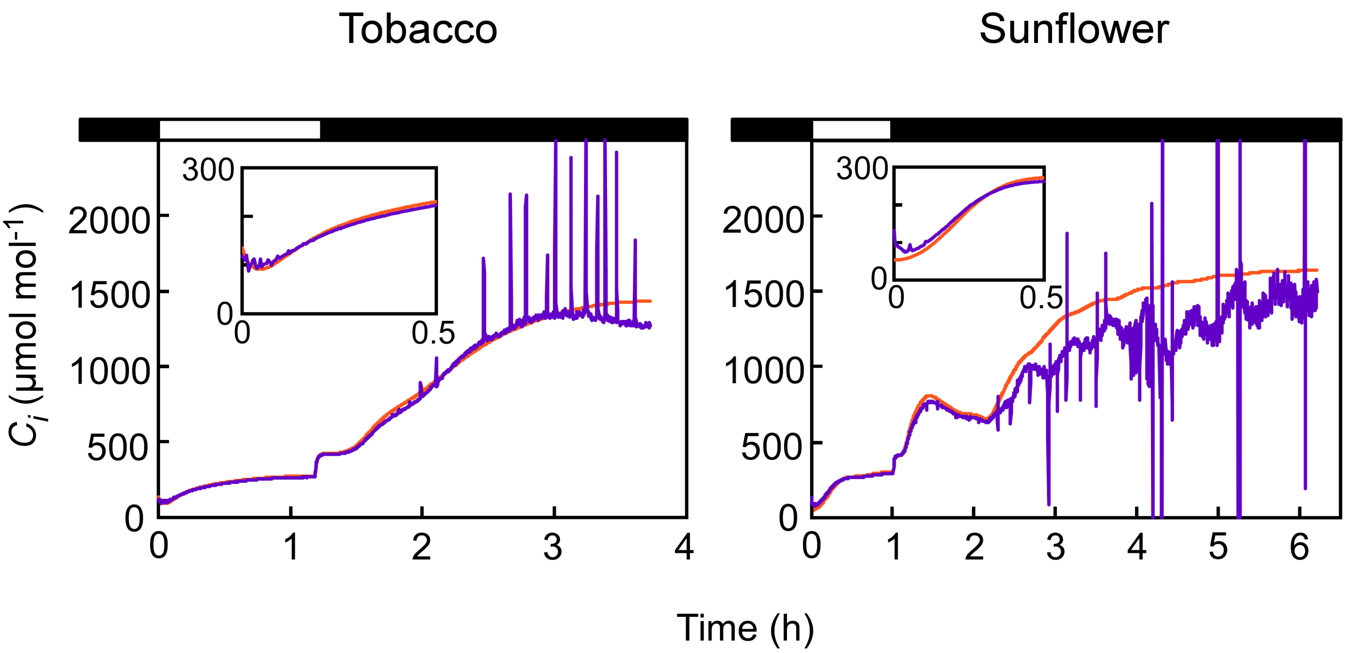


**Fig. S1.** Effect of intercellular $CO_2$ gradients on the $C_{i(c)}$. Data for Fig. 1 is shown. The $C_{i(c)}$ was corrected for $CO_2$ gradients after the correction for cuticle conductance ($C_{i(c),cut+grad}$, purple line) with the mean $g_{ias}$ determined anatomically; 617 mmol $CO_2$ $m^{-2}$ $s^{-1}$ for tobacco and 538 mmol $CO_2$ $m^{-2}$ $s^{-1}$ for sunflower, respectively (MS#1). Note the absence of change in the $C_{i(c),cut+grad}$ from the $C_{i(c),cut}$ (Fig. 3).
