## Supplementary material for "Minimum conductance in leaves—cuticle, leaky stomata, or water vapor saturation?": Fig. S2

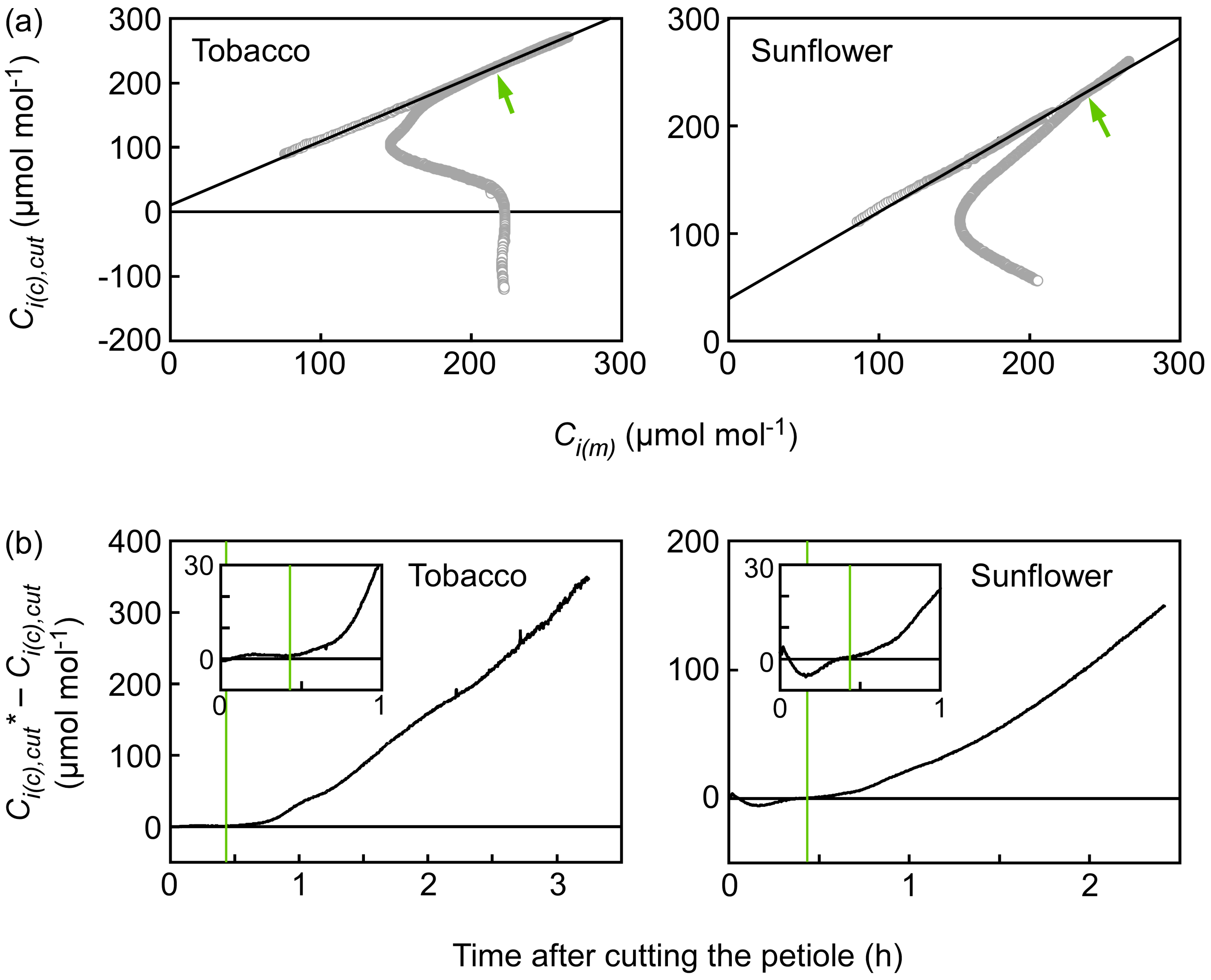


**Fig. S2.** Initiation of unsaturation in leaf airspace. Relationships between $C_{i(c),cut}$ and $C_{i(m)}$ throughout the experiments for Fig. 6 (a), and difference of the $C_{i(c),cut}$ expected from the $C_{i(m)}$ for the attached leaves ($C_{i(c),cut}{}^{*}$) and the $C_{i(c)}$ observed after the leaf detachment (b). In (a), linear function was fit throughout the maximum $C_{i(m)}$ [$Y$=0.988$X$ + 10.7 ($R^2$=0.9995) for tobacco and $Y$=0.807$X$ + 39.6 ($R^2$=0.9967) for sunflower, respectively], and were used for calculating the $C_{i(c),cut}{}^{*}$ in (b). Green arrows and lines indicate when the $C_{i(c),cut}$ started to consistently decrease from the $C_{i(c),cut}{}^{*}$.
