## Supplementary material for "Minimum conductance in leaves—cuticle, leaky stomata, or water vapor saturation?": Fig. S3

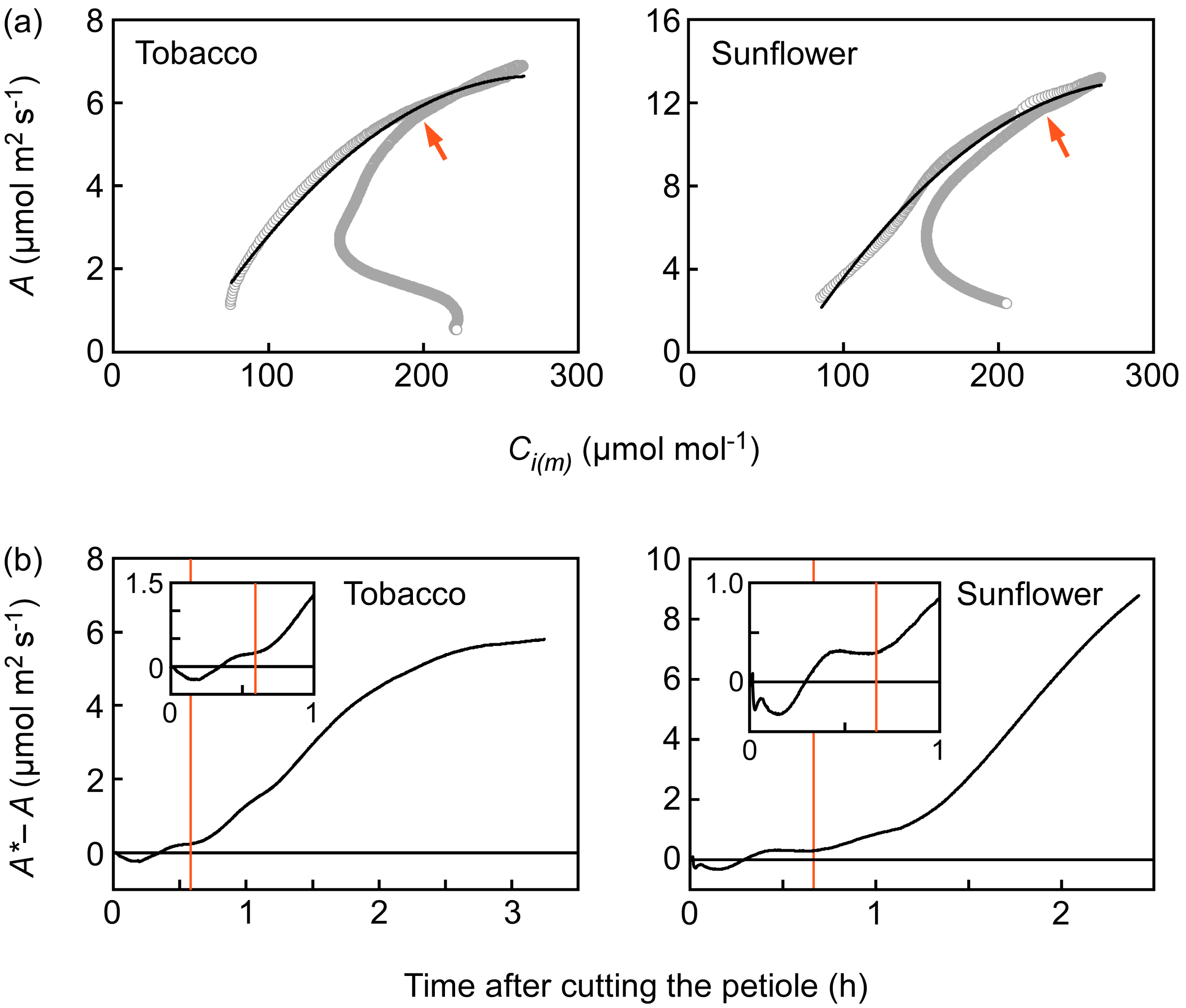


**Fig. S3.** Initiation of suppression in photosynthetic activity. Relationships between $A$ and $C_{i(m)}$ throughout the experiments for Fig. 6 (a), and difference of the $A$ expected from the $C_{i(m)}$ for the attached leaves ($A^*$) and the $A$ observed after the leaf detachment (b). In (a), quadratic curve was fit throughout the maximum $C_{i(m)}$ [$Y$=-1.26•10$^{-4}$$X^2$ + 6.93•10$^{-2}$$X$ -2.87 ($R^2$=0.9931) for tobacco and $Y$=-2.50•10$^{-4}$$X^2$ + 14.7•10$^{-2}$$X$ - 8.67 ($R^2$=0.9878) for sunflower, respectively], and were used for calculating the $A^*$ in (b). Orange arrows and lines indicate when the $A$ started to consistently decrease from the $A^*$.
