## Supplementary material for "Minimum conductance in leaves—cuticle, leaky stomata, or water vapor saturation?": Fig. S4

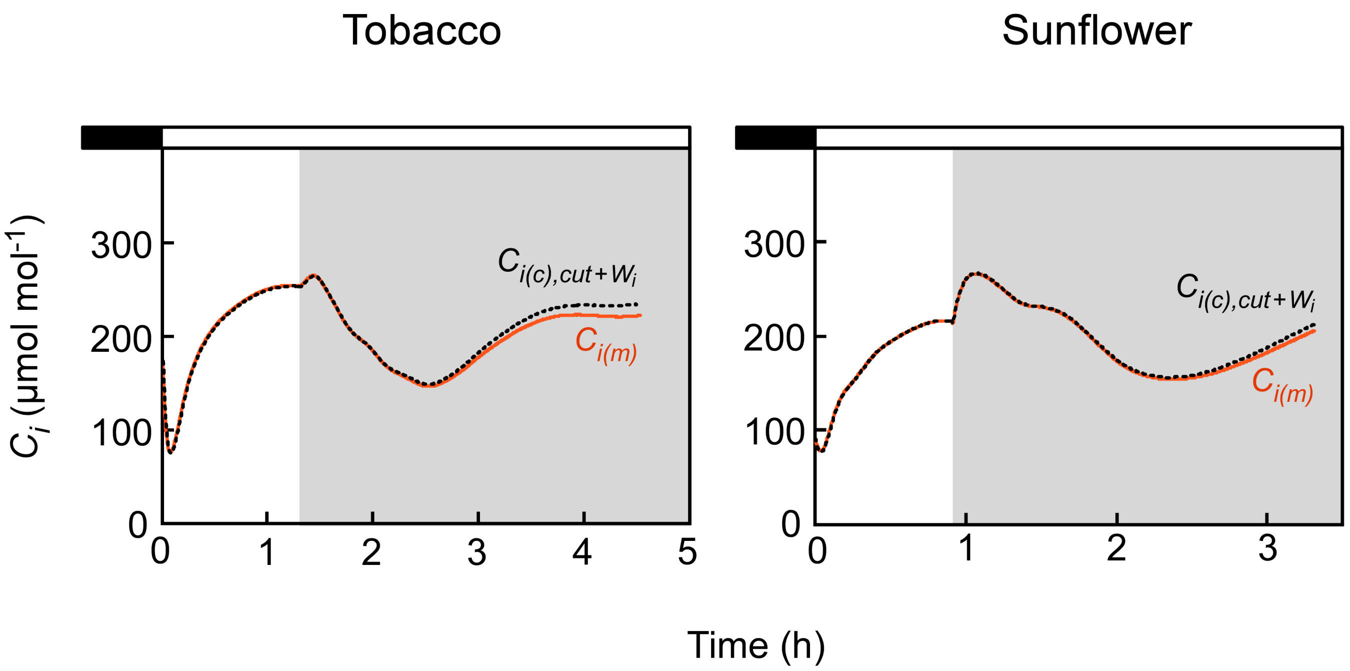


**Fig. S4.** Correction for water vapor concentration in leaf airspace ($W_i$). Data for Fig. 6c are shown. The $C_{i(c),cut}$ was corrected ($C_{i(c),cut+Wi}$, black dot line) with the unsaturated $W_i$ ($W_i$').
