## Supplementary material for "Minimum conductance in leaves—cuticle, leaky stomata, or water vapor saturation?": Fig. S5

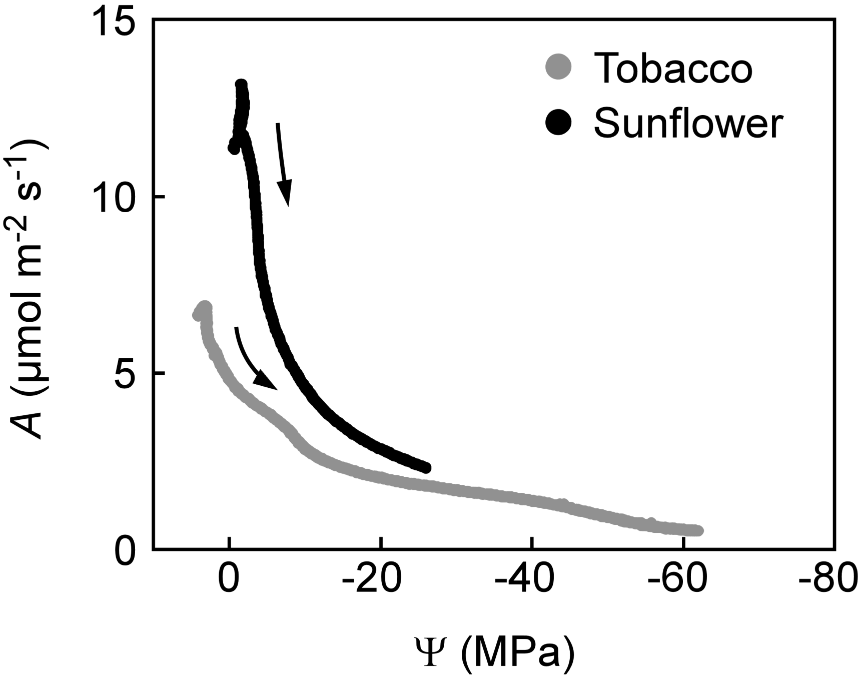


**Fig. S5.** Relationships between *A* and water potential (Ψ) of evaporating surface in leaves after cutting the petiole. Data for Fig. 6 are shown. Arrows indicate time progress.
